# A Computational Framework for the Administration of 5-Aminovulinic Acid before Glioblastoma Surgery

**DOI:** 10.1101/2023.12.07.570672

**Authors:** Jia Zeng, Nicholas J. Moore

## Abstract

5-Aminolevulinic Acid (5-ALA) is the only fluorophore approved by the FDA as an intraoperative optical imaging agent for fluorescence-guided surgery in patients with glioblastoma. The dosing regimen is based on rodent tests where a maximum signal occurs around 6 hours after drug administration. Here, we construct a computational framework to simulate the transport of 5-ALA through the stomach, blood, and brain, and the subsequent conversion to the florescent agent protoporphyrin IX at the tumor site. The framework combines compartmental models with spatially-resolved partial differential equations, enabling one to address questions regarding quantity and timing of 5-ALA administration before surgery. Numerical tests in two spatial dimensions indicate that, for tumors exceeding the detectable threshold, the time to peak flouresent concentration is 2-7 hours, broadly consistent with the current surgical guidelines. Moreover, the framework enables one to examine the specific effects of tumor size and location on the required dose and timing of 5-ALA administration before glioblastoma surgery.

## I. INTRODUCTION

Glioblastoma is one of the most aggressive types of cancer that starts within the brain. It has been categorized as a grade IV astrocytic lesion by the World Health Organization. The median survival time for glioblastoma is only 15 months [2, 19]. Currently, the best treatments for patients with glioblastoma is maximal resection. However, it is often difficult to distinguish tumor from normal neighboring brain using only traditional white-light microscopy. As such, fluorescence-guided surgery (FGS) has been developed to help surgeons overcome this challenge [18, 28].

The first use of fluorescence imaging in surgery dates back to 1948 when surgeons injected fluorescein intravenously to locate neoplastic cells [15]. Since then, a number of fluorophores have been used in clinical practice. Among them, 5-ALA is the only fluorophore approved by the FDA as an intraoperative optical imaging agent in patients with suspected high grade glioma. 5-ALA was firstly introduced to FGS by Dr. Walter Stummer in 1998 [26]. It is a naturally occuring metabolite for hemoglobin biosythesis. 5-ALA itself is non-fluorescent, but the lack of the enzyme ferrochelatase in tumor cells sharply decreases the synthesis rate of hemoglobin, causing the heme precursor protoporphyrin IX (PpIX) to accumulate. The PpIX accumulated inside the tumor radiates red fluorescence when illuminated with blue light [7].

Current computational models for 5-ALA dosing were designed for skin cancer treatment, where 5-ALA is applied topically as opposed to orally [1, 22, 24]. The dosing regimen for administration of 5-ALA for FGS, an oral dose of 5-ALA HCl solution at 20 mg/kg body weight given 3 hours in advance [6], is based on rodent tests, where a fluorescence peak was seen 6 hours after the injection [23]. To give enough time for anesthesia, monitoring and performing the craniotomy, it was decided that the patient should take the medication 3 hours before surgery to allow for the maximal PpIX fluorescence during tumor resection.

The main question considered in this work is: What is the time required for the concentration of fluorophore (PpIX) inside the tumor to reach a maximum so that surgery can be performed during under optimal conditions? We address this question with a computational framework that accounts for the transport of 5-ALA from the stomach, through the bloodstream, to the brain, and the subsequent conversion to PpIX at the tumor site. Compartmental models based on ordinary differential equations (ODEs) are sufficient to describe the transport of 5-ALA through the stomach and the bloodstream [10], but the assumption of rapid mixing underlying these models does not hold inside the brain, where 5-ALA slowly diffuses inwards from the blood-brain barrier. We therefore resolve the spatial dependence of 5-ALA diffusion within the brain and its subsequent conversion to PpIX through numerical solution of partial differential equations (PDEs). The ODE and PDE systems are coupled through boundary conditions at the blood-brain barrier that control the transfer rate of 5-ALA across that interface. A 2D implementation of this model yields predictions for the peak florescence times that are broadly consistent with available data, such as the aforementioned rodent tests. Moreover, the framework enables one to examine the specific effects of tumor size and location on the peak florescence time and on the minimal dose required for florescence.

## II. NUMERICAL METHODS

In this section, we introduce the computational framework used to simulate the 5-ALA and PpIX dynamics. Section II A introduces the general model for 5-ALA dynamics within the stomach, blood, and brain, where the brain is represented by an arbitrary 2D or 3D geometry. Section II B specializes this model to a simple 2D semi-circular brain geometry and describes the finite-difference methods employed. II C treats the conversion of 5-ALA to PpIX and the subsequent transport inside the tumor using stochastic particle methods.

### A. General framework for 5-ALA dynamics

In order to be used in glioblastoma surgery, 5-ALA must travel from the stomach to the bloodstream and then across the blood-brain barrier. Once inside the brain, 5-ALA converts to PpIX at the tumor site. The rapid mixing within the stomach and bloodstream enables the use of simple ODE-based compartmental models. Letting *c*_1_(*t*) and *c*_2_(*t*) denote the concentration of 5-ALA within the stomach and bloodstream respectively (see Table I for the list of symbols used), the ODEs governing the transport of 5-ALA within these two regions are given by [24]:

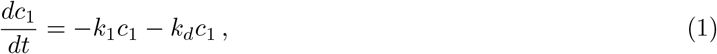

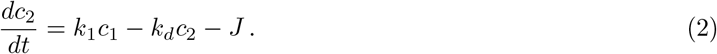

**TABLE I:**
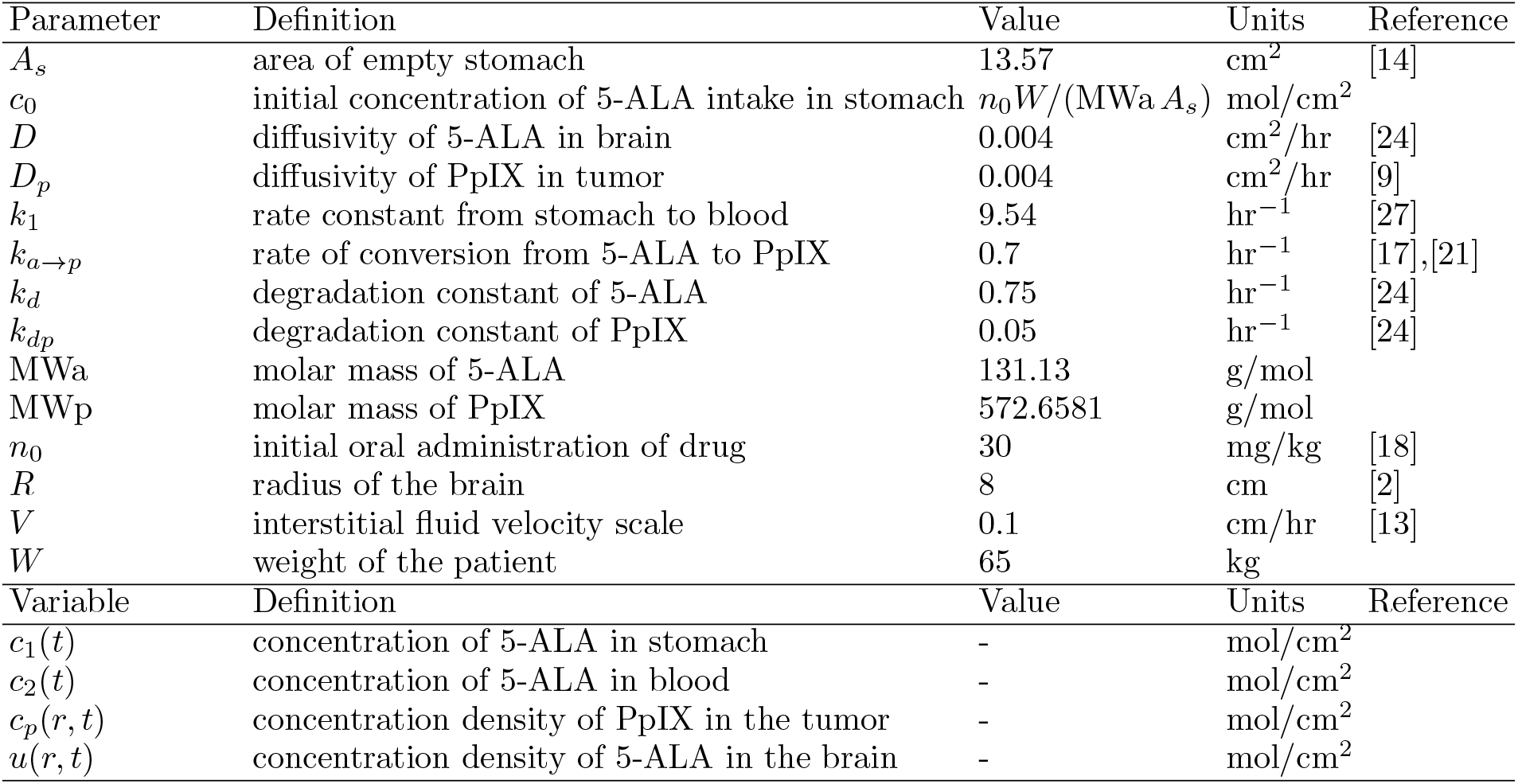
Notation for physical parameters and variables used.

In these ODEs, the first term on each right-hand-side, 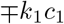, represents transport of 5-ALA from the stomach to the blood with rate parameter *k*_1_. The second term on each right-hand-side represents degradation of 5-ALA in the stomach and bloodstream respectively with rate parameter *k*_*d*_. Equation (1) is decoupled from the rest of the system and can be immediately solved as

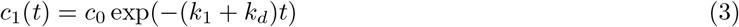

where the initial 5-ALA concentration in the stomach *c*_0_ is given in terms of other parameters, as shown in Table I. The last term in Eq. (2), denoted *J*, represents the flux of 5-ALA across the blood-brain barrier. Importantly, the value of *J* is unknown *a priori* and will be determined self-consistently by coupling to the PDE-model for transport within the brain.

We first consider a brain represented by an arbitrary domain Ω lying in two or three spatial dimensions and with boundary *∂*Ω. Transport of 5-ALA within the brain is results from diffusion and degradation as governed by the PDE:

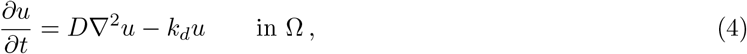

where *u* is the concentration of 5-ALA, taken to be *u* = 0 at the initial time. 5-ALA enters the brain along the section of boundary *∂*Ω_*b*_ representing the blood-brain barrier. At this interface, the concentration of 5-ALA must be continuous, giving the Dirichlet boundary condition:

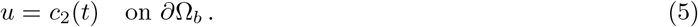

There may exist another section of the boundary *∂*Ω_*i*_ that is impervious to 5-ALA, giving the Neumann condition

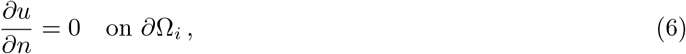

where *n* is the direction normal to *∂*Ω_*i*_.

At this stage, *c*_2_(*t*) is unknown due to the presence of the unknown flux *J* in Eq. (2). Hence, the system consisting of Eqs. (1)–(2) and (4)–(6) is currently underdetermined. The resolution is that the flux of 5-ALA exiting the bloodstream relates to the flux of 5-ALA entering the brain. Accounting for this relationship closes the system of equations so that a unique solution can be found.

To this end, let *c*_3_ represent the spatially-averaged concentration of 5-ALA within the brain,

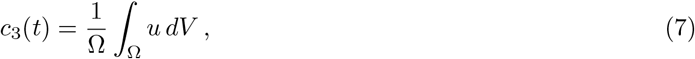

where we use the short-hand Ω to denote the volume (or area in 2D) of domain Ω. Then computing the rate of change of *c*_3_(*t*), using the PDE (4), applying the divergence theorem, and simplifying gives

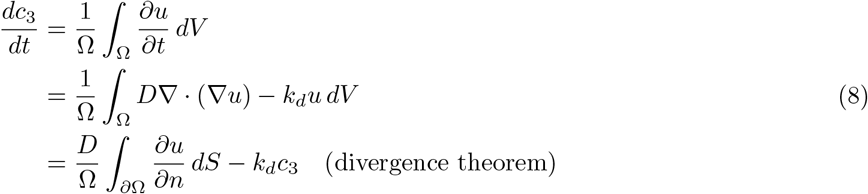

On the final line, the last term represents degradation of 5-ALA while the first term identifies the flux *J* of 5-ALA crossing the blood-brain barrier as

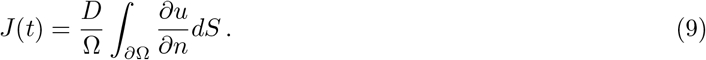

With *J* expressed in terms of boundary data of the main unknown *u*, the system consisting of Eqs. (1)–(2), (4)–(6), and (9) comprises a closed system of equations.

### B. 2D Implementation of 5-ALA dynamics

In this paper, we consider a simple 2D representation of the brain, namely a semicircular domain with radius *R* as illustrated in Fig. 1. We will use polar coordinates (*r, θ*) with the semicircle represented by 0 *< r < R* and 0 *< θ < π*. The perimeter represents the blood-brain barrier, giving boundary condition

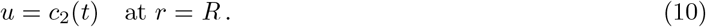

**FIG. 1:**
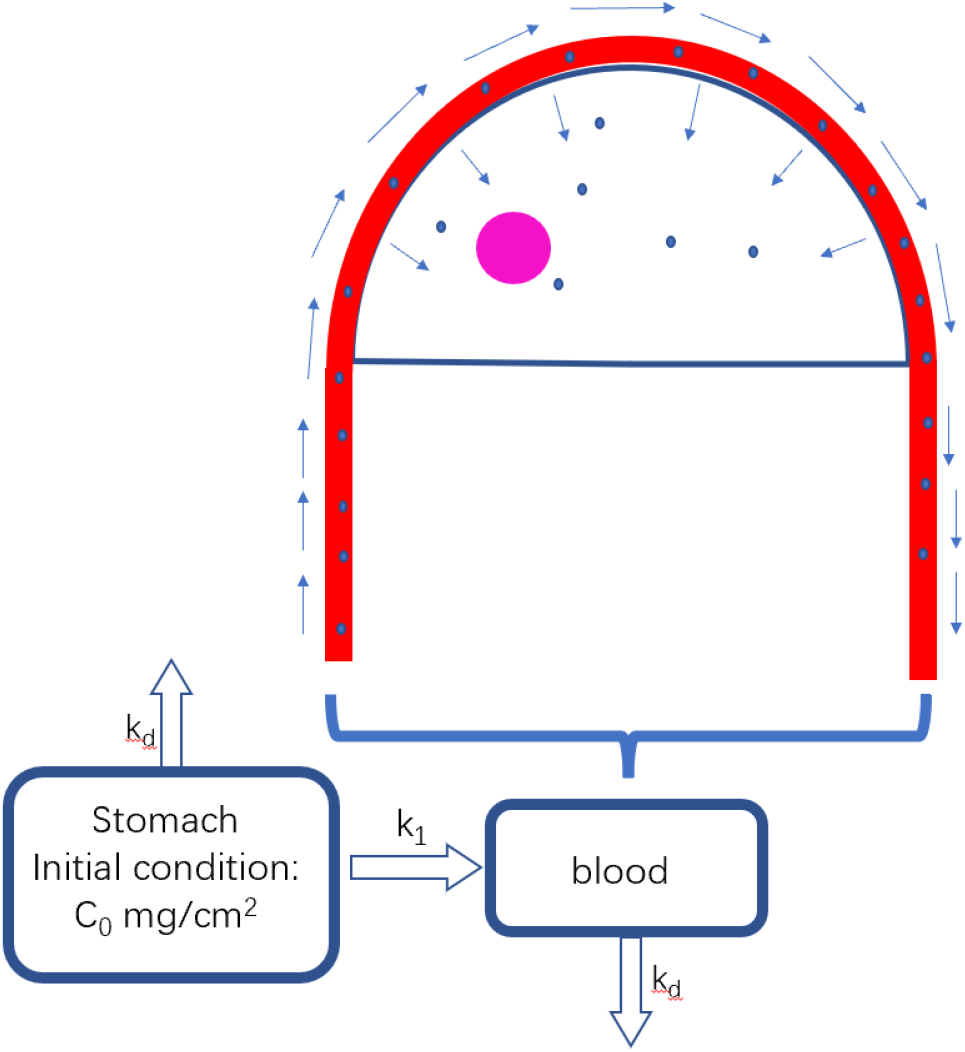
Shematic illustration of the transport of 5-ALA (blue particles) through the blood (red domain), the brain (semicircular domain) and the tumor site (purple circle).

Meanwhile, the baseline represents an impervious surface, giving Neumann conditions:

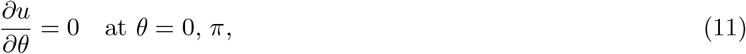

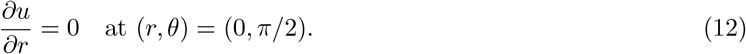

Due to the symmetry present in these boundary conditions and in the initial condition, the solution must remain rotationally invariant for all times; that is, the solution *u* = *u*(*r, t*) is independent of *θ*. Using Eq. (9) with domain area Ω = *πR*^2^*/*2 and boundary conditions (10)–(12), the flux of 5-ALA across the blood-brain barrier becomes

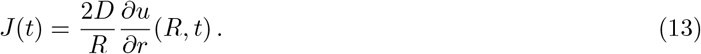

Hence, for the simple semicircular geometry, the coupled system of equations for unknowns *c*_2_(*t*) and *u*(*r, t*) can be summarized as

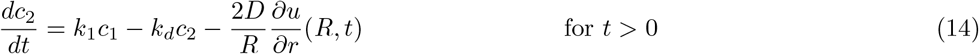

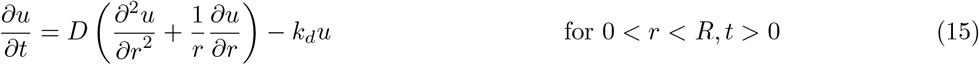

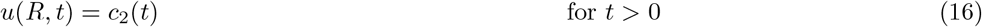

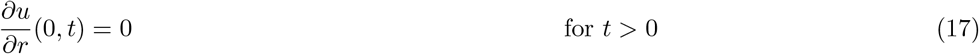

with vanishing initial conditions, *c*_2_(0) = 0 and *u*(*r*, 0) = 0, and with *c*_1_(*t*) given by Eq. (3).

We solve the coupled ODE-PDE system (14)–(17) numerically using centered finite differences and explicit time stepping. In particular, suppose the domain 0 *< r < R* is divided into *N*_*r*_ equal sub-intervals, each of width Δ*r* = *R/N*_*r*_, so that grid points are given by *r*_*j*_ = *j*Δ*r* for *j* = 0, 1, · · · *N*_*r*_. Meanwhile, time is discretized as *t*_*n*_ = *n*Δ*t* where Δ*t* is the step size. Let subscripts and superscripts indicate the spatial and temporal discretization respectively, so that 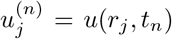 and 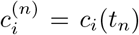 for *i* = 1, 2. Applying second-order centered-differences in space and the forward Euler method in time gives the following update rules:

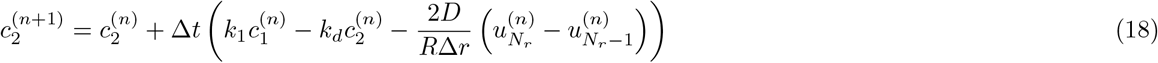

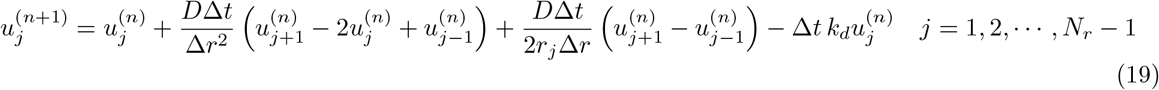

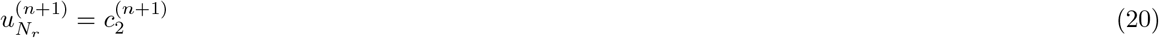

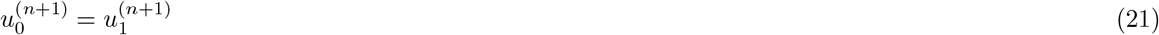

The penultimate equation represents the outer Dirichlet boundary condition (16) while the final equation represents the Neumann boundary condition at the origin (17). Importantly, the time step is chosen to satisfy the Courant–Friedrichs–Lewy (CFL) condition for numerical stability, Δ*t ≤* Δ*r*^2^*/*(2*D*) [4]. In numerical experiments discussed below we set Δ*r* = 0.1 cm and Δ*t* = 0.005 hr.

### C. Generation and transport of Protoporphyrin IX

Upon reaching the tumor site, 5-ALA converts to PpIX through a chain of chemical reactions [8, 22]. In the absence of diffusion and other transport processes, the conversion from 5-ALA to PpIX can be described as a first-order reaction,

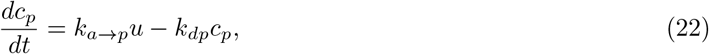

where *c*_*p*_ represents the concentration of PpIX, *k*_*a*→*p*_ is the conversion rate, and *k*_*dp*_ is the degradation rate of PpIX. Transport processes, however, modify Eq. (22) to take the form of an advection-diffusion PDE [3, 12, 13]

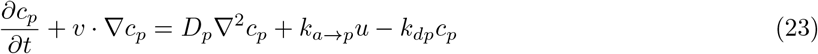

In the above, the spatially dependent PpIX concentration, *c*_*p*_(*x, y, t*), is advected by interstitial velocity *v* = (*v*_*x*_, *v*_*y*_) and diffuses with rate *D*_*p*_, in addition to the generation and degradation processes.

Since the tumor is located arbitrarily in the brain, the concentration of PpIX does not exhibit any exploitable symmetry in the polar-coordinate system as 5-ALA did. We will therefore treat the PpIX evolution in Cartesian coordinates *c*_*p*_(*x, y, t*) for simplicity, and we will solve Eq. (23) numerically in Langevin form using a stochastic method. Compared to a finite-difference method in two spatial dimensions, the stochastic method sacrifices some accuracy but offers advantages in a low (dimension-independent) computational cost, better numerical stability, and relative ease of implementation. It is also worth noting that it is trivial to generalize the stochastic method to 3D.

In Langevin form, the continuous PpIX concentration field *c*_*p*_(*x, y, t*) is represented by a finite number of computational particles. Each particle does not necessarily represent an individual PpIX molecule, but rather a specified mass of PpIX residing within a small control volume. In particular, we set the number of computational particles per mol of PpIX to be *N*_*m*_ = 10^12^.

Over the course of the simulation, computational particles are created due to the conversion of 5-ALA to PpIX and destroyed due to the degredation of PpIX. Specifically, new particles are created within each computational cell lying in the tumor. For grid point (*r*_*i*_, *θ*_*j*_), consider the surrounding computational cell [*r*_*i*_ − Δ*r/*2, *r*_*i*_ + Δ*r/*2] × [*θ*_*j*_ − Δ*θ/*2, *θ*_*j*_ + Δ*θ/*2], where Δ*r* = *R/N*_*r*_ as before and we use the same number of grid points in the *θ*-direction, Δ*θ* = *π/N*_*r*_. The area of cell (*i, j*) is Δ*A*_*i*_ = *r*_*i*_Δ*r*Δ*θ*. Focusing first only on the conversion of 5-ALA to PpIX in (22), the expected number of new PpIX particles generated within cell (*i, j*) over time Δ*t* is given by

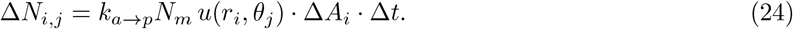

Degredation will be treated separately. Above, the value Δ*N*_*i,j*_ may be non-integer. Therefore, at each time step, we first generate 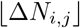 new particles in each cell, where 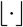 is the floor function. Newly generated particles are placed randomly in the cell using a uniform distribution. Then, to represent the remainder of Δ*N*_*i,j*_, we generate one last particle with probability 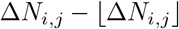, thus guaranteeing that the expected number of new particles is Δ*N*_*i,j*_. Finally, to represent degradation, we allow each particle to be destroyed with probability 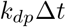.

Once generated, the computational PpIX particles are transported by advection and diffusion according to Eq. (23). In Langevin form, the Cartesian coordinates (*X*(*t*), *Y* (*t*)) of a particle evolve according to the stochastic ODE [11]:

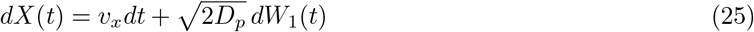

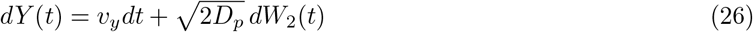

where (*dW*_1_, *dW*_2_) represents two-dimensional Brownian motion (in differential form). That is, *dW*_1_ and *dW*_2_ are independent, identically distributed random numbers selected from a normal distribution with mean zero and variance *dt*:

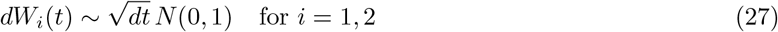

When PpIX approaches the tumor boundary, the pressure gradient existing between healthy tissue and the tumor drives PpIX radially outwards [13]. This process gives rise to an interstitial velocity that exists within a narrow region surrounding the tumor boundary. We take the width of the region to be 0.05 cm. For a computational PpIX particle within this region, the interstitial velocity takes the form [13]

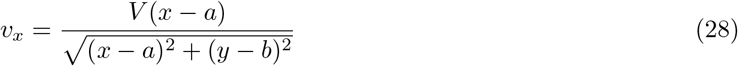

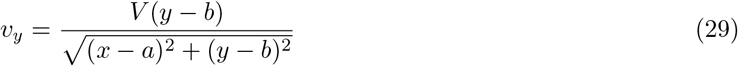

where (*a, b*) represents the tumor center and *V* = 0.1 cm/hr is a velocity scale. For particles farther than 0.05 cm from the tumor boundary, we take *v* = 0.

## III. RESULTS AND DISCUSSION

We now use the numerical methods introduced in Section II to simulate the coupled dynamics of 5-ALA and PpIX within a stomach-blood-brain system represented by the simple 2D geometry shown in Fig. 1. Ultimately, we seek to address the question of what is the optimal timing for 5-ALA administration before FGS.

### A. Diffusion of 5-ALA in the brain

We first simulate the transport of 5-ALA by solving the coupled ODE-PDE system (14)–(17) with numerical method given by Eqs. (18)–(21). Table I lists the values of the control parameters used. In particular, we simulate the diffusion of 5-ALA through a semi-circular brain of radius *R* = 8 cm. Recall that, due to the symmetry of the domain and boundary conditions, the 5-ALA distribution remains independent of *θ* for all time. Figure 2(a) shows the radial dependence of 5-ALA as it varies with time. At early times, 5-ALA is concentrated predominantly near the blood-brain boundary, *r* = 8 cm, as seen by the dark blue curves. As time passes, 5-ALA diffuses inwards while also decreasing in overall amount due to degradation (light blue and green curves). Notice that, due to the combination of inward diffusion and overall degradation, the time of peak 5-ALA concentration depends on location. For example, for *r* = 7.8 cm, the 5-ALA concentration peaks at roughly 2.5 hours. For smaller *r*, the peak occurs later, while for larger *r* the peak occurs earlier.

**FIG. 2:**
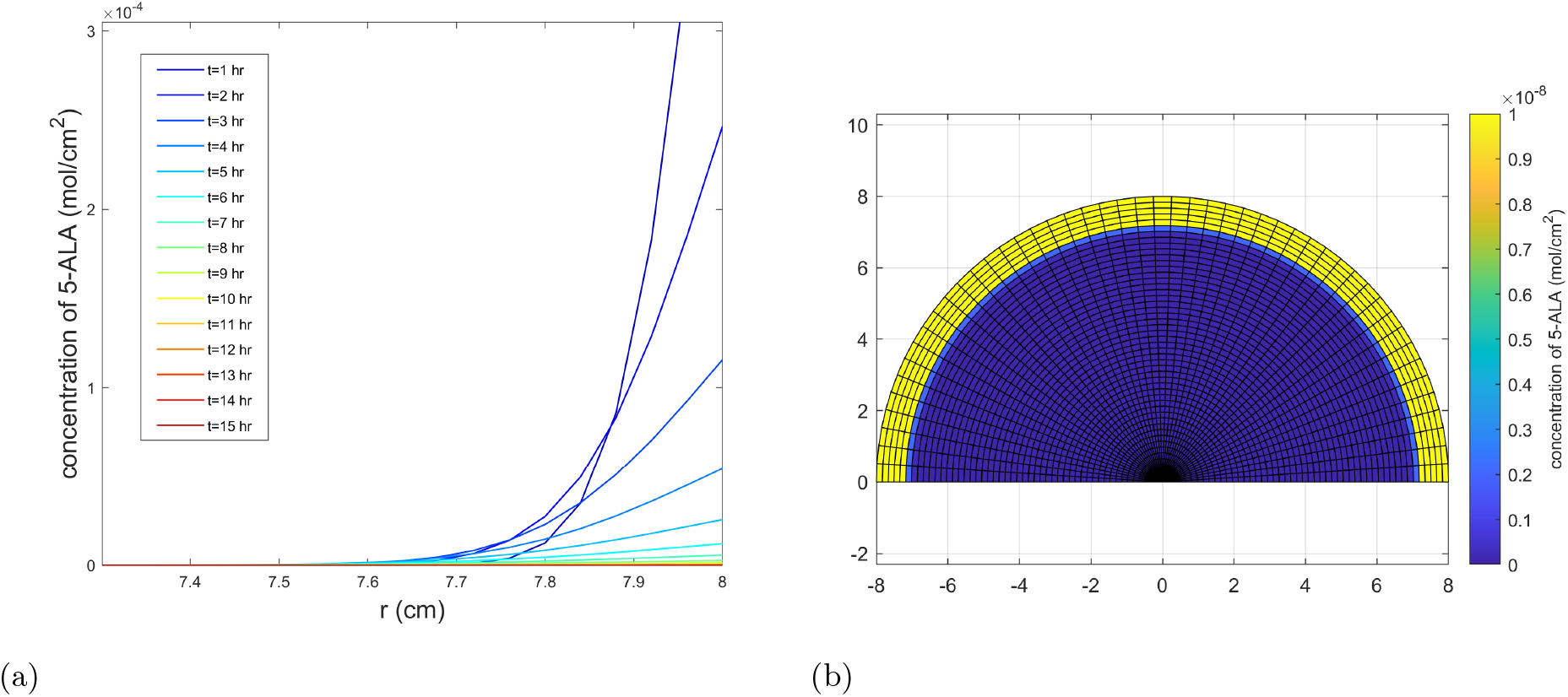
a) Radial distribution of 5-ALA as it varies with time. The 5-ALA diffuses from the boundary *r* = *R* inwards while also degrading. b) Visualization of the 5-ALA distribution in the semi-circular brain at a time of *t* = 5 hours after dosing. At this time, 5-ALA has diffused a distance of roughly 1 cm, forming a diffusive layer around the outer boundary.

In the model, the 5-ALA actually diffuses through a semi-circular domain, and Fig. 2(b) provides a visualization of this process. The figure shows the distribution of 5-ALA at time *t* = 5 hours after ingestion, demonstrating the 5-ALA to diffuse from the outer boundary towards the center. By this time, a diffusive layer of roughly 1 cm has formed around the outer boundary. Inwards of this diffusive layer, the 5-ALA concentration remains close to zero.

### B. Generation of PpIX

We next consider the generation and transport of PpIX within a tumor. We consider a circular tumor located arbitrarily (i.e. with no particular symmetry) inside the semicircular brain, as depicted in Fig. 3(a). In this figure, a tumor of radius *R* = 2 cm is centered at (*x, y*) = (4, 4). This arrangement results in a gap of roughly 0.34 cm between the tumor boundary and brain boundary. Later, we also consider other values of tumor radius and position not illustrated in the figure.

**FIG. 3:**
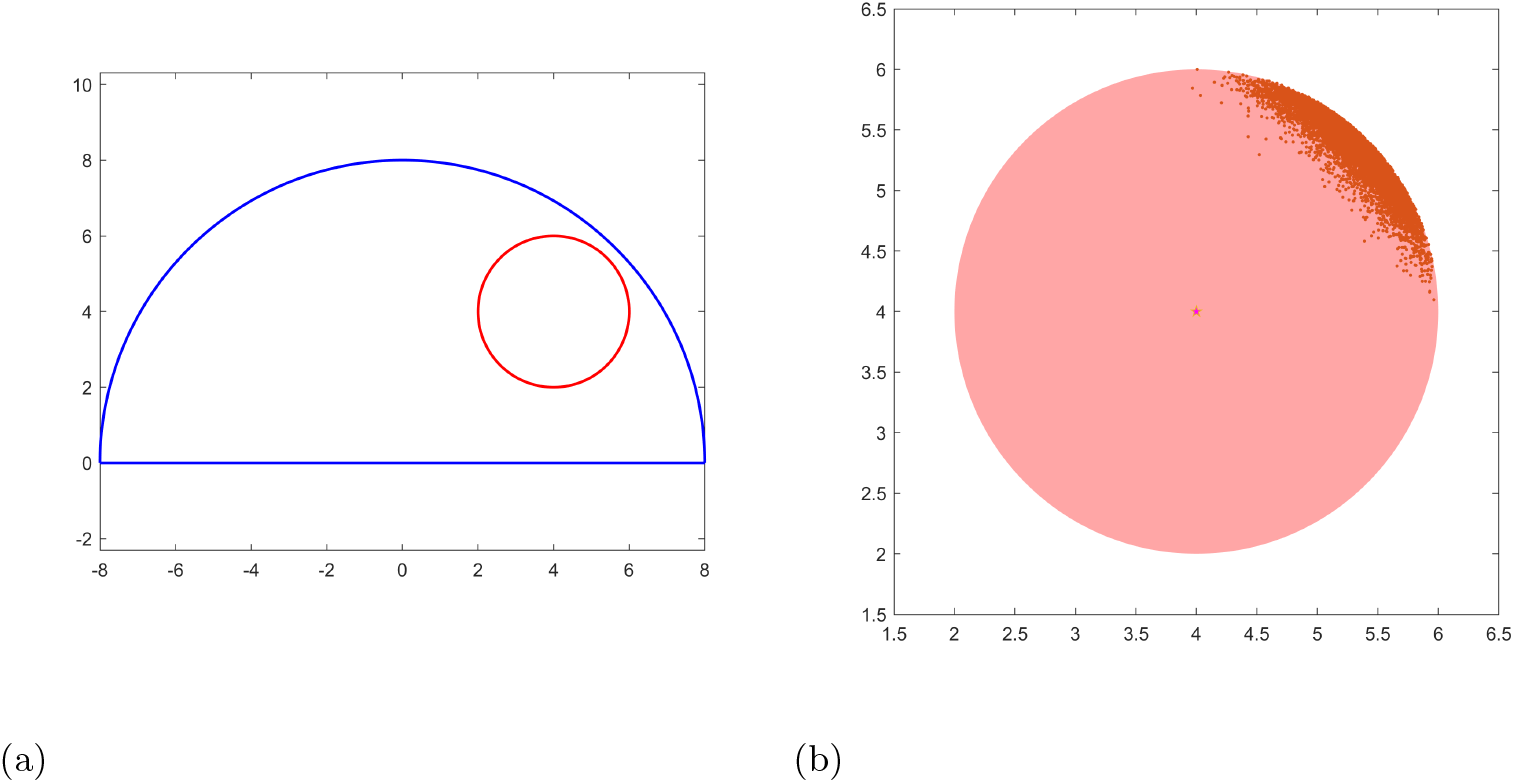
(a) Tumor of radius 2 cm centered at (*x, y*) = (4, 4) within a semi-circular brain of radius *R* = 8 cm. (b) The particulate distribution of PpIX for the tumor shown in (a) at time *t* = 5.6 hours corresponding to the peak of the PpIX concentration. The PpIX is concentrated in the upper-right of the tumor, nearest the brain boundary where the 5-ALA concentration is highest.

The PpIX dynamics described by Eq. (23) are simulated using the stochastic particle method outlined by Eqs. (24)–(26). In particular, PpIX is generated proportional to the local amount of 5-ALA, and it is transported by a combination of diffusion and interstitial velocity. The resulting PpIX concentration field is represented by a finite number of point particles, as illustrated in Fig. 3(b). This figure shows the PpIX particle field for the tumor depicted in Fig. 3(a) at the time of peak PpIX concentration (*t* = 5.6 hours). Notice the PpIX field is concentrated in the upper-right sector of the tumor, that is, the sector that lies closest to the outer brain boundary, *r* = 8 cm, where the 5-ALA concentration is highest. The distribution in Fig. 3(b) is therefore consistent with expected qualitative behavior, since PpIX generation is proportional to the local 5-ALA concentration. Even though the PpIX is concentrated in only a small region, its florescence would illuminate the entire tumor, so long as the PpIX concentration exceeds the detection threshold.

### C. Timing of 5-ALA administration for FGS

We now consider the main question underlying this work: What is the optimal timing for 5-ALA administration before FGS? Since the PpIX concentration controls the amount of fluorescence during FGS, we will use the PpIX distributions of the type shown in Fig. 3(b) to estimate the corresponding PpIX concentration. In particular, we integrate the PpIX field over the entire tumor and then divide by the estimated area occupied by the PpIX particles. This procedure provides an estimate for the maximum PpIX concentration in the tumor at any given time.

Figure 4 shows the estimated PpIX concentration as it varies with time for three cases of tumor size and location. First, Fig. 4(a) shows the case of a 1.5-cm-radius tumor centered at (4, 4), implying a gap of 0.84 cm between the tumor edge and brain boundary. As seen here, the PpIX concentration increases slowly at early times, peaks at roughly *t* = 10 hours, and then decays. Notice the PpIX concentration remains below 0.5 ng/cm^2^ at all times, implying a very low PpIX concentration overall. Figure 4(b) shows the data for a tumor with the same center (4, 4) but a radius of 2 cm, giving a gap of 0.34 cm. The trend is qualitatively similar, but the peak occurs earlier (*t* = 5.6 hours) with a much higher value (600 ng/cm^2^) as compared to (a). Here, the smaller gap between the tumor edge and brain boundary exposes the tumor to higher 5-ALA concentrations, resulting in much higher PpIX concentrations.

**FIG. 4:**
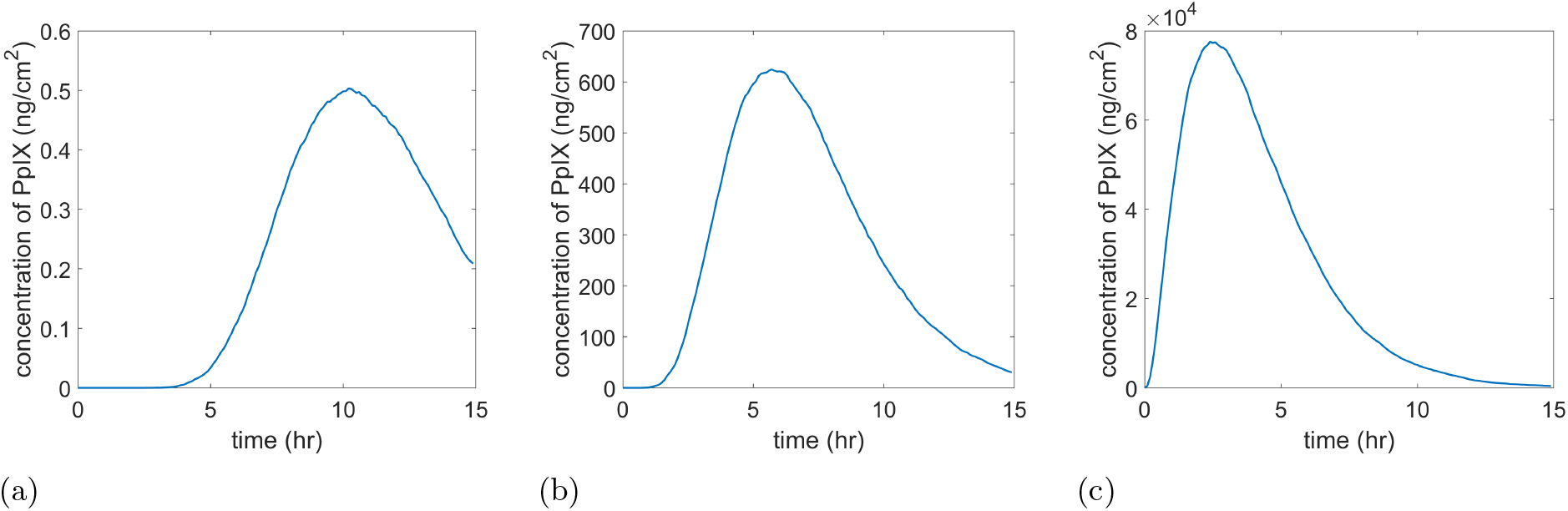
Maximum concentration of PpIX inside the tumor as it varies with time. (a) 1.5-cm-radius tumor centered at (4, 4). (b) 2-cm-radius tumor centered at (4, 4). (c) 2-cm-radius tumor at 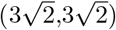 so that the edge of the tumor contacts the brain boundary.

Figure 4(c) shows the same data for an extreme case of a 2-cm-radius tumor with center 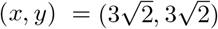, implying that the edge of the tumor lies tangent to the brain boundary; i.e. the gap between the two is nonexistent. This arrangement exposes the tumor to the highest 5-ALA concentrations possible. As seen in Fig. 4(c), the resulting PpIX concentration is much greater in magnitude, with a peak concentration of 7.5 × 10^4^ ng/cm^2^ occurring at *t* = 2.5 hours.

As seen from this data, small changes in the size and location of the tumor can lead to moderate changes in the timing of peak PpIX concentration and enormous changes in the value of that peak concentration. Not all of these cases, however, lead to concentrations that would be detectable during FGS. The detection threshold of PpIX during FGS has been estimated to be on the order of 10 ng/ml [16]. To relate this estimate to our 2D simulations, we multiply by a length scale representing a characteristic depth. In particular, we choose a typical tumor diameter, 4 cm, although other length scales, such as the tumor radius or the brain diameter/radius, could be chosen. With this conversion, we estimate 40 ng/cm^2^ as the PpIX detection threshold for our 2D simulations. Returning to Fig. 4, the case of Fig. 4(a) lies two orders of magnitude below the detectable threshold, while the cases of Fig. 4(b) and (c) are well above the threshold. Thus, while the 1.5-cm tumor represented in Fig. 4(a) implies an optimal waiting time of roughly 10 hours between 5-ALA ingestion and surgery, this prediction is irrelevant since the PpIX would not fluoresce sufficiently. On the other hand, the 2-cm tumor cases shown in Figs. 4(b) and (c), with waiting times of 2-6 hours, are relevant for FGS. We note that FGS is typically performed for high-grade gliomas [5] with diameters of at least 4 cm [25], which corroborates our numerical findings that larger tumors are more easily detectable during FGS.

While Fig. 4 showcases a handful of possible tumor sizes and locations, we explore the influence of tumor location more systematically in Fig. 5. Here, we consider a tumor of radius 2 cm with the distance to the brain boundary varied. The figure shows the ideal time, that is the time of peak PpIX concentration, on the left axis. The right axis shows the value of the peak concentration on a log scale. As expected, increasing the distance between the tumor center and brain boundary increases the ideal waiting time and reduces the peak concentration. While the ideal time varies over a moderate range of 2–12 hours, the peak concentration value varies over 6 orders of magnitude. This peak concentration value is substantially more sensitive to tumor position than is the waiting time. The horizontal line in the figure shows our estimated detection threshold of 40 ng/cm^2^. As seen in the figure, the maximum PpIX concentration exceeds this threshold in cases in which the distance is smaller than 2.5 cm, or equivalently the gap between the tumor edge and brain boundary is smaller than 0.5 cm. Tumors positioned farther from the brain-blood barrier would not be detectable during FGS and therefore the waiting times of 8–12 hours are irrelevant. The detectable tumors, positioned closer to the blood-brain tumors, exhibit ideal waiting times in the range of 2–7 hours, which is broadly consistent with waiting times employed in FGS [7].

**FIG. 5:**
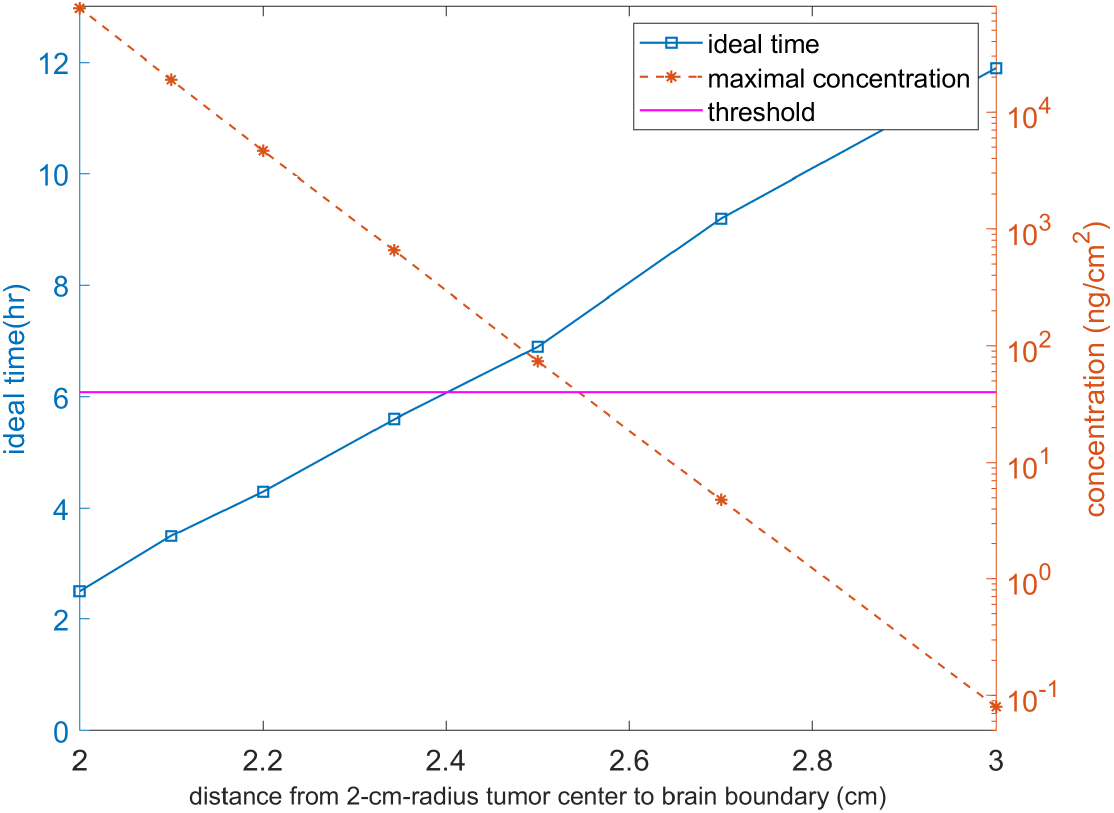
Impact of tumor location on FGS variables. As the location of a 2-cm-radius tumor is varied systematically, the time and value of the peak PpIX concentration is measured. The time of the peak (left axis) increases with distance to the brain boundary, while the peak concentration (right axis) decreases with distance. The peak concentration value, shown on a log scale, varies over 6 orders of magnitude, while the ideal time varies in the range of 2–12 hours. The horizontal line shows the estimated detection threshold of 40 ng/cm^2^. The cases that exceed the threshold for detectability exhibit ideal waiting times in the range 2–7 hours, consistent with waiting times used in FGS.

## IV. CONCLUSION AND FUTURE WORK

This paper introduces a computational framework to simulate the chemical and physical processes underlying 5-ALA-based fluorescence-guided glioblastoma surgery. The framework couples ODE compartment models for 5-ALA transport within the stomach/blood to a spatially-resolved PDE description of 5-ALA transport within the brain and the subsequent generation of PpIX. Importantly, the transport of 5-ALA across the blood-brain barrier is treated as an unknown that is determined self-consistently through solution of the coupled ODE-PDE system. We employ a finite-difference method for the 5-ALA transport and a stochastic particle method for the PpIX generation and transport. Fundamentally, the framework applies to either two or three spatial dimensions. As a proof of concept, we provide in this paper 2D results on how the time and value of peak florescence depends on tumor size and position.

Even though the implementation discussed here is two dimensional, the model appears to capture the essential physical processes of 5-ALA/PpIX dynamics and proves to be broadly consistent with FGS guidelines employed in practice. In particular, for tumors exceeding the detectability threshold, numerical tests suggest a peak PpIX time of 2-7 hours, consistent with the 6-hour peak time observed in rodent tests [23], and consistent with the current dosing regimen of a 3-hour lead time between 5-ALA administration and the beginning of surgery [6]. Furthermore, numerical tests indicate that tumors substantially smaller than 4 cm in diameter would be difficult to detect during FGS unless they happen to be positioned very near the blood-brain barrier. This finding is consistent with the current guidelines of using FGS for tumors at least 4 cm in diameter [5, 25]. Tumors are known to grow in size over timescales of weeks to months [20]. While our model shows that this growth would increase the detectability of the tumor, the growth is also known to induce severe effects such as intracranial pressure and edema [20], and hence chemotherapy is advisable for smaller tumors that are not located sufficiently near an artery [19].

In the numerical implementation given here, the brain is modeled as a 2D semi-circular domain with the outer perimeter representing the blood-brain barrier. In reality, the blood-brain barrier is comprised of a complex network of arteries that penetrate into the brain. Future implementations could account for this complex network of arteries to achieve greater realism. Further extensions could account for the full, three-dimensional geometry of the brain and arteries, while also accounting for different diffusivity values of white and gray matter in the brain. Moreover, controlled laboratory experiments could help determine more accurate parameter values involved in 5-ALA and PpIX transport (e.g. the digestion parameter *k*_1_ and the diffusivity *D* in the brain).

With such improvements, we envision that this computational model could be used to address important diagnostic questions leading up to glioblastoma surgery. For example, after measuring a tumor’s size and location with a CT or MRI scan, the model could determine, with high accuracy, answers to the following questions:

1. Will the tumor be detectable during FGS?
2. What is the minimum quantity of 5-ALA required to detect the tumor in order to avoid cytotoxicity from overdosing?
3. What is the optimal time the patient should intake 5-ALA before surgery?

The 2D implementation presented here provides a proof of concept and suggests that the accuracy required for clinical application could be realistically achieved by a 3D implementation. By changing certain parameters, the framework may also apply to nanoparticle delivery in tissue generally, which could be particularly useful in time-sensitive scenarios.

